# Human DND1-RRM2 forms a non-canonical domain swapped dimer

**DOI:** 10.1101/2020.03.05.978023

**Authors:** Pooja Kumari, Neel Sarovar Bhavesh

## Abstract

Human DND1 (Dead end protein homolog1) is an RNA binding protein. DND1 plays pivotal role in animal development and has been implicated in cancer. DND1 consists of two RNA recognition motifs (RRMs) in tandem and a double stranded RNA binding domain at the C terminal separated by 40 residues flexible linker. The conserved RNP site in the RRM1 domain helps in specific RNA recognition while the RNP sites in RRM2 are not well conserved. DND1 has been reported to be involved in inhibition of microRNA access to target mRNA and it also associate with CCR4-NOT complex that targets mRNA. In order to understand this intriguing contrasting molecular function, we have determined the 2.3 Å resolution crystal structure of the human DND1 RRM2 domain. The structure revealed an interesting non-canonical RRM fold that is maintained by the formation of a domain swapped dimer between β_1_ and β_4_ strands across two chains. The domain swapping is attributed by a hinge loop between α_2_ and β_4_ that helps in mediating a domain swap forming anti-parallel β sheets. We have delineated the structural basis of stable dimer formation using the residue level dynamics of protein explored by NMR spectroscopy and MD simulations. Our structural and dynamics studies demonstrate the molecular basis for the dimerization of the RRM2 domain and shed light on the possibility for this motif for interaction with other proteins which helps in transcription regulation.

**Graphical Abstract:** 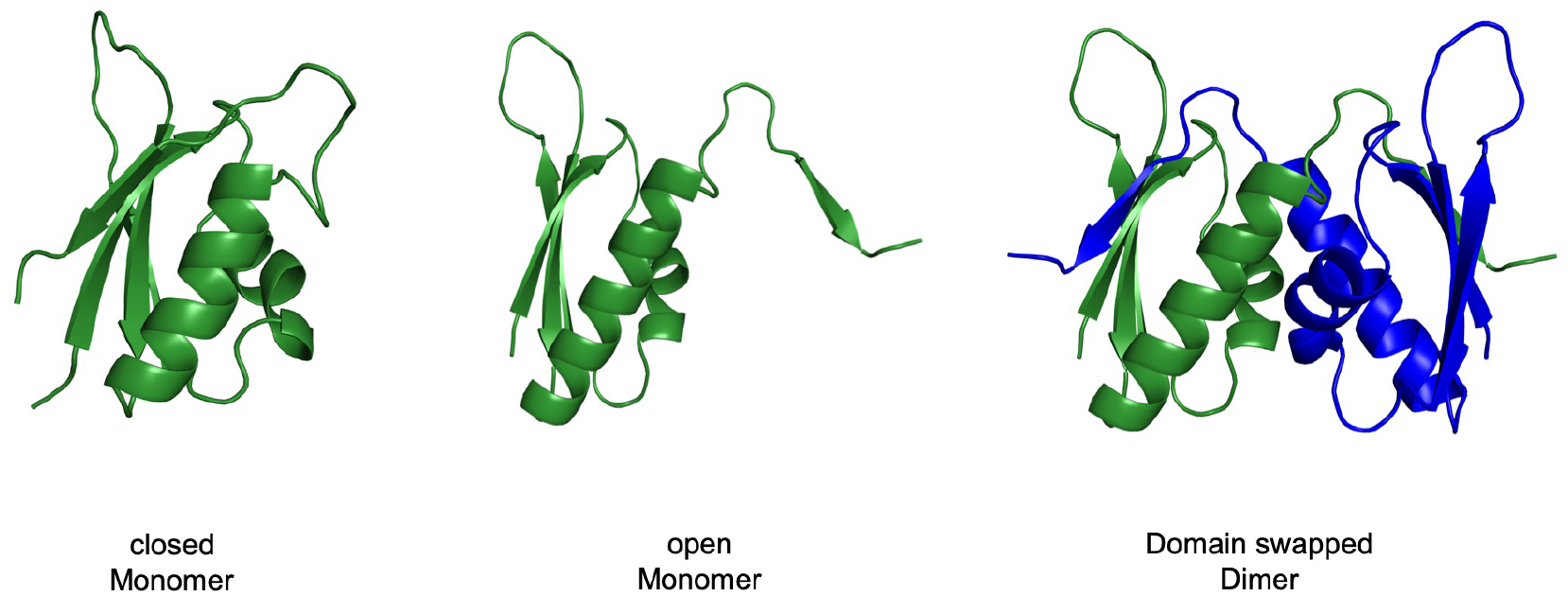

**Highlights:**

- First report of a domain swapped dimer formation by an RRM identified through crystal structure determination of the human DND1 RRM2 domain.
- RRM2 exhibit domain swapped dimerization attributed by hinge loop and disulfide bond formation.
- Dimer formation is under redox regulation.
- Major determinants of swapping were identified.
- DND1 RRM2 is not involved in RNA recognition.

## Introduction

Human DND1 (Dead end protein homolog1) also known as DND MicroRNA-Mediated Repression Inhibitor 1 is an RNA binding protein containing two RRMs (RNA recognition motif), RRM1, RRM2 domains, in tandem and a double stranded RNA binding domain at the C-terminal separated by 40 residues flexible linker. Located at position q31.3 on human chromosome 5, it is highly conserved in vertebrates and is essential for migration and viability of primordial germ cells in zebrafish (Weidinger et al., 2003). Complete loss of DND1 protein in mice leads to early embryonic lethality (Zechel et al., 2013). Introduction of a nonsense mutations (R190X) named as Ter in RRM2 of DND1 in mice and (W289X) in dsRBD of rats, results in formation of a truncated DND1 protein, which is followed by germ cell depletion, found to be oncogenic and result in testicular germ cell tumors (Noguchi and Noguchi, 1985; Northrup et al., 2012; Youngren et al., 2005). In humans, reports of uncontrolled regulation at protein level or mutation in the gene have been associated with testicular cancer, tongue squamous cell carcinoma (Linger et al., 2008; Liu et al., 2010; Sijmons et al., 2010).

The molecular function studied so far associated with DND1 has been set forth from studies in zebrafish, rodents and human cells. There is a disparity in the molecular function so far unraveled. DND1 is known to positively modulate the stability and translation of AU rich element containing mRNA. It has been also reported to associate with proteins and macromolecular complexes, which destabilize mRNAs. Studies done on human cells and zebrafish germ cells show that DND1 protein interacts with the 3’-UTR (untranslated region) of mRNAs and inhibits miRNA regulated repression of complementary target mRNAs. On the basis of experimental observation in which sequence in the vicinity of miRNA targets were fairly conserved. It has been proposed that DND1 interaction to mRNA obstructs the accessibility of miRNA/AGO complexes on 3′ untranslated region (UTR) of p27, LATS2, NANOG, LIN28 mRNA in zebrafish and human (Kedde et al., 2007; Zhu et al., 2011). On the contrary, using PAR-CLIP and co-immunoprecipitation studies Yamaji et al. have recently proposed that DND1 not only binds to UU(A/U) trinucleotide motif on the 3′ untranslated regions of mRNA but also interacts with proteins from the CCR4 complex. CCR4-NOT comprises of deadenylating enzymes responsible for mRNA decay and DND1 interaction was mapped to the N terminal of CNOT1 scaffold unit (Yamaji et al., 2017). The specification of transcripts for deadenylation is usually mediated through adapter proteins that interact with the enzyme complexes as well as its cognate RNA sequences (Webster et al., 2019). Although no information on the three-dimensional structure is available, structure prediction tools utilizing sequence-based prediction methods propose multi-domain architectures for DND1 protein shows 3 separated domains (Waterhouse et al., 2018).

Here, we report the crystal structure of the DND1 RRM2 domain at 2.3 Å resolution which reveals the formation of a domain swapped dimer. Using a combination of NMR spectroscopy and MD simulation we further deduce mechanism of the conformational flexibility in the hinge loop. The conformational switch is under redox regulation, shedding light on its varied function.

## Results

### RRM2 of DND1 contains pseudo RNP motifs

DND1 consists of three domains as predicted by DELTA-BLAST (Domain Enhanced Lookup Time Accelerated BLAST) which are RRM1 (58-136), RRM2 domain (139-217) and dsRBD (254-332) double stranded RNA binding domain. Analysis of DND1 sequence showed significant bias consisting 50% residues which are aliphatic having distinctly high content of leucine, alanine, glycine and proline, 7% aromatic, 20% neutral, 13% basic and 7% acidic residues (Figure S1 A). To identify the structural features and their conservation, which are responsible for DND1 function, DND1 sequence was aligned from various vertebrate organisms shown in figure S1-B. We observed that RRM1 was very well conserved with its Ribonucleoprotein (RNP) motifs, RNP1 and RNP2, responsible for specific RNA interaction, corresponding with RNP1 and RNP2 consensus sequence which is (R/K)-G-(F/Y)-(G/A)-(F/Y)-V-X-(F/Y) and (L/I)-(F/Y)-(V/I)-X-(N/G)-L, respectively (Burd and Dreyfuss, 1994). However, major differences were found in RRM2 domain and in dsRBD at the C-terminal region, where the RNP motifs were neither well defined nor well conserved. PAR-CLIP studies performed by Yamaji et al showed that DND1 predominantly binds to UU(A/U) trinucleotide motif in the 3′ untranslated regions of mRNA (Yamaji et al., 2017). We investigated the binding using Isothermal titration calorimetry and titrated RRM2 with UUUUUU and UUAUUU RNA sequences and found that RRM2 shows no interaction to these RNA sequences which is shown in figure S1 C-D. As the RNP sites containing the aromatic residues, which normally interact with RNA, are not well conserved. Hence, RRM2 does not play a role in RNA recognition.

### DND1-RRM2 forms a stable domain swapped dimer

The DND1 RRM2 (139-217) domain contains leucine and proline rich sequence and does not bear any sequence homology when searched on protein data bank with known structure. This intrigued us to determine the structure and gain mechanistic insights about its regulation inside the cell. We over-expressed the DND1-RRM2 domain construct in *Escherichia coli* Bl21 (*DE3*) Codon plus cells and purified to homogeneity as our efforts to obtain both RRM1 and RRM2 domain together did not succeed. This RRM2 construct henceforth numbered from 1 to 100 aa contained a 21 amino acid long N-terminal hexa-histidine tag M1-M21 with thrombin cleavable site followed by the RRM2 domain E22-K100 residues. The domain architecture is shown in figure 1A. The purified DND1 RRM2 domain crystallized and the native crystal diffracted X-rays to a resolution of 2.4 Å. DND1 RRM2 domain lacked homology to any known protein structure expect with RBM47 (PDB ID 2DIS) calculated using NMR spectroscopy. Therefore, the crystal structure of DND1 RRM2 was determined using selenium based single wavelength anomalous diffraction (SAD) method. The crystal diffracted to 2.3Å and the belonged to I4_1_ space group. DND1 RRM2 domain crystallized with one molecule in the asymmetric unit and four copies in the unit cell. The molecule in the asymmetric unit had continuous electron density from N- to C-terminal except for the 12 residues comprising the N-terminal hexa-histidine tag. Data processing and structure refinement statistics is given in table 1.

**Table 1.** Data processing and structure refinement statistics, Numbers in parentheses denote the highest-resolution shell.

|  |  |
| --- | --- |
| Space Group: | I 4<sub>1</sub> |
| Unit Cell Parameters (Å, °) | $a=44.62, b=44.62, c=111.33$ $\alpha=\beta=\gamma=90$ |
| Matthews coefficient (Å<sup>3</sup>Da<sup>-1</sup>) | 2.64 |
| Solvent Content (%) | 53.41 |
| Data-collection temperature (K) | 100 |
| Detector | Pilatus 6M |
| Wavelength(Å) | 0.972 |
| Resolution limits (Å): | 31.57 - 2.3 (2.34-2.30) |
| Total reflections | 29990 (1616) |
| Unique reflections | 4476 (237) |
| Data completeness % | 92.2 (100) |
| R<sub>merge</sub> | 0.092 (0.52) |
| I/$\sigma$ I | 15.7 (3.7) |
| Anomalous completeness | 89.4 (95.2) |
| Multiplicity | 6.7 (6.8) |
| Anomalous multiplicity | 3.5 (3.6) |
| CC<sub>1/2</sub> | 0.997 (0.921) |
| Wilson B-factor (Å<sup>2</sup>) | 32.0 |
| Mosaicity (°) | 0.2 |
| **Refinement statistics** |  |
| R-Value Work: | 0.21 |
| R-Value Free: | 0.24 |
| B factor | 34.0 |
| **Ramachandran Statistics** |  |
| Preferred region | 98.85 % |
| Allowed region | 1.15 % |
| outliers | 0 |
| **Data Deposition** |  |
| PDB id | 6LE1 |

**Figure 1.**
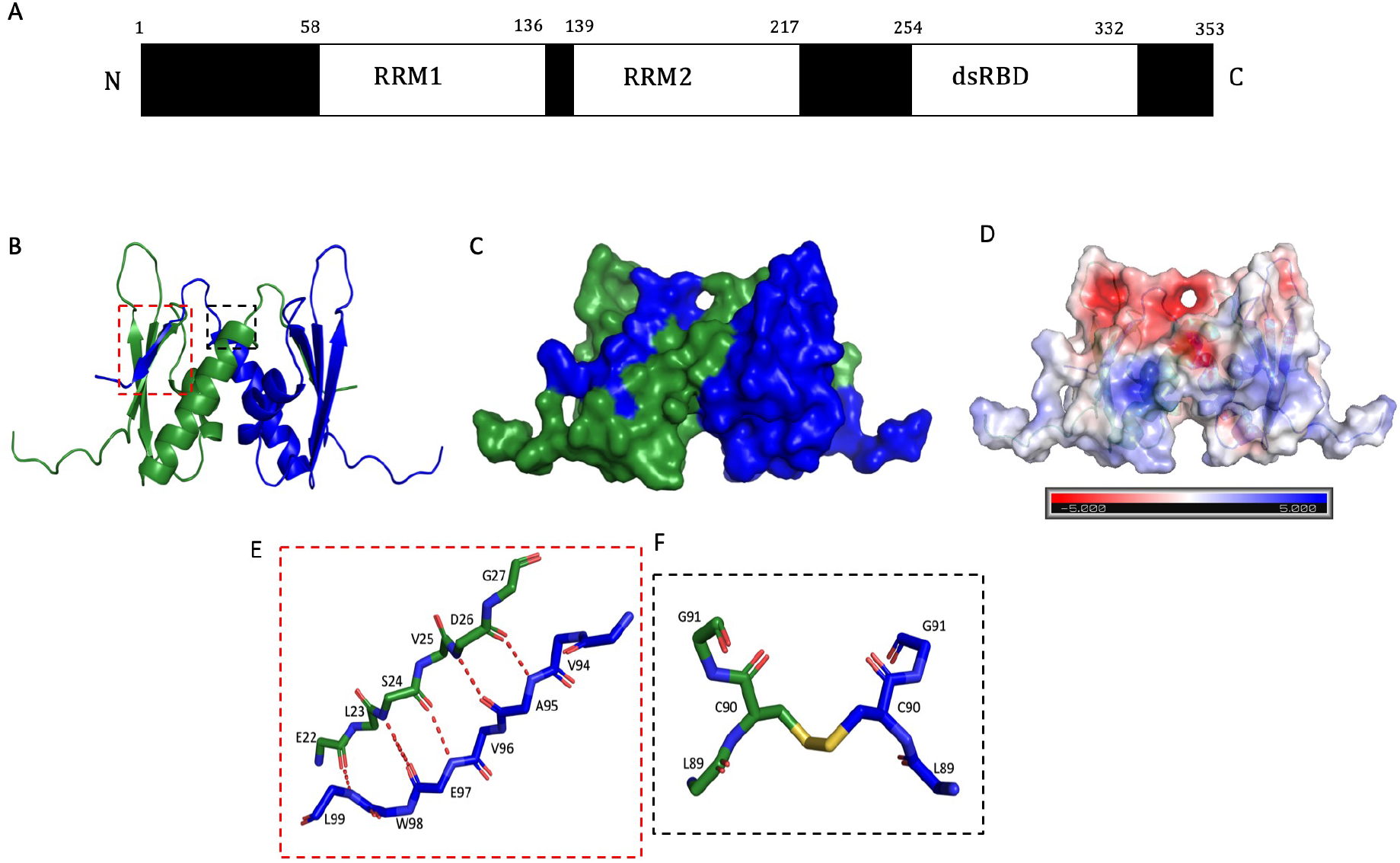
Domain organization and crystal structure of DND1 RRM2 dimer. **(A)** Domain organization of DND1 protein showing its domains and their boundaries. **(B)** Overall structure in ribbon representation showing 2 chains in blue and green showing domain swapped dimer formation. **(C)** Space-filling model of the DND1-RRM2 domain-swapped dimer is shown to highlight the quaternary arrangement of the monomers **(D)** Electrostatic potential surface generated by the APBS method as implemented in PyMOL Molecular Graphics System, Version 2.0 Schrödinger, LLC showing charge polarity. **(E)** The backbone trace of the exchanged region is shown, and the hydrogen bonds formed between the two chains are shown by red dashed lines. **(F)** Disulphide bond between two cysteine residues from chain A and chain B shown by yellow solid line.

Structural analysis revealed a striking non-canonical feature where complete RRM motif comprising of four β sheets and two α helices forming a β_1_-α_1_-β_2_-β_3_-α_2_-β_4_ fold was maintained by formation of a globular dimer structure (figure 1B-C). A domain swapped homodimer between β_1_-strand of one monomer and β_4_-strand of another monomer of DND1 RRM2 resulted in formation of RRM fold. We refer to each RRM chain with the chain labels A and B. A hinge loop between α_2_-helix and β_4_-strand mediates a domain swap forming an anti-parallel β-sheet between the chains. The solvent accessible area and buried surface area for the protein is 7255.7 Å^2^ and 2000.6 Å^2^ with a solvation energy of −55.2 kcal/mol. Domain-swapped dimerization is one way by which loop hydrophobic residues which are surface exposed can be stabilized. The dimer assembly comprising of two RRM2 molecules had a surface area of 10989.8 Å^2^ and buried area of 4330.9 Å^2^ and free energy of solvation gained upon the formation of the domain swapped dimer structures is −16.6 Kcal/mol. Examination of surface electrostatics potential shown in fig 1D revealed a bias in polarity with strong negative charge near the apical loops while strong positive charge near the basal loop regions. The domain swapped anti-parallel β-sheet formation is stabilized by hydrogen bonds between E22 to D26 in β_1_ and A95 to L99 in β_4_ and disulphide bond between C90 between chain A and chain B shown in figure 1E-F. Complete list of all the interactions and their distance between the two chains are given in table S1.

### Monomer dimer switch under redox regulation

Given the dimeric domain organization of the protein in the crystal, we set out to define whether this dimeric association holds true in solution. We further concentrated the purified protein and ran it again on size exclusion chromatography column calibrated using five standard proteins and found two peaks corresponding to monomer as well as dimer population. Therefore, the RRM2 dimer association observed in the crystal was also present in solution, instead of it being a crystallization artifact. The dimer is further stabilized by formation of a disulphide bond utilizing the single cysteine residue C90 between the two chains. We set out to define weather this dimer formation is under redox regulation. For this we added 5mM β-merceptoethanol to the buffer solution containing DND1 RRM2. We found that condition in which we used 5mM β-merceptoethanol showed only monomer population with absorbance unit of 1400 mAU (figure 2A) while in its absence the protein showed two populations comprising of monomer as well as dimer with lower absorbance unit of 450 mAU (figure 2B). We also validated this result using non-reducing SDS-PAGE where the loading dye lacked β-merceptoethanol, the monomer and dimer could be clearly distinguished as seen in figure 2C. Hence, the monomer to dimer switch is also under redox regulation, as only monomer population was observed in reducing condition even at high concentration.

**Figure 2.**
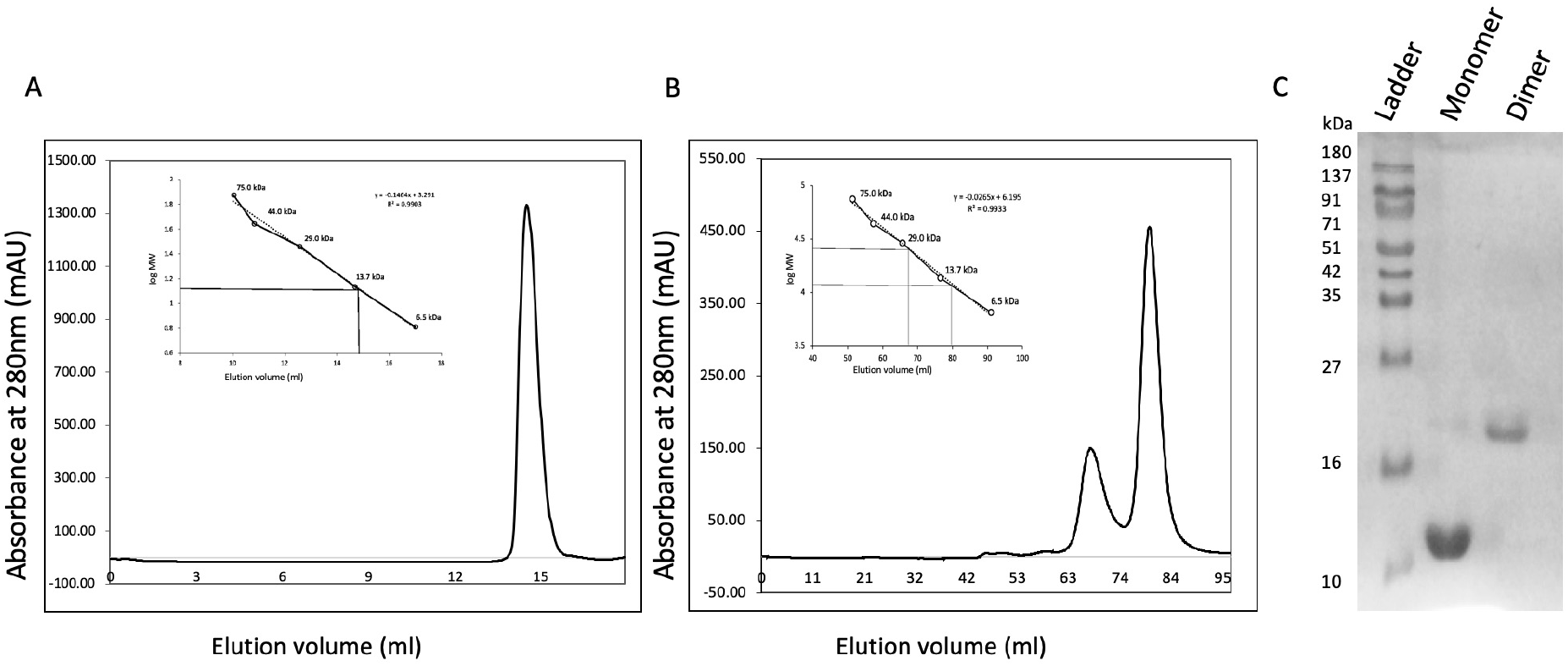
Monomer dimer switch under redox regulation. **(A)** SEC profile of DND1 RRM2 in presence of 5mM β-merceptoethanol showing only monomer. **(B)** SEC profile of DND1 RRM2 in absence of β-merceptoethanol showing both monomer and dimer population in solution. **(C)** Non-reducing SDS-PAGE showing the protein ladder in lane 1, monomer in lane 2 and dimer in lane 3.

The binding sites on proteins plays an important role in determining its interactions be it protein-protein, protein nucleotide, protein-ligand or protein-drug and hence it is very important for predicting protein function and gives rationale for drug designing. Schrödinger SiteMap (Schrödinger, LLC) tool allowed us to gain insights of regions within the binding site which are suitable for occupancy by ligand, hydrogen-bond donors, acceptors, hydrophobic groups or metal-binding functionality (Halgren, 2007, 2009). This allowed us to distinguish the sub-regions of different binding site and classify the drugability of a protein. We analyzed the protein for binding site identification using Sitemap in the Schrödinger suite and we found that the loop region in the apical and basal surface can serve as a binding site, which in turn could be responsible for protein-protein interaction (figure S1 E-F). Also, in the case of dimer the four α-helices from the two RRM molecules could serve as a potential binding site having a hydrophobic core with a drugability score of 1.37 indicating that it is a druggable site shown in figure S1 G.

The closest structural homology entry for DND1-RRM2 chain A from an exhaustive search of the Protein Data Bank (PDB) using DALI (Holm and Rosenström, 2010) shows proteins which are involved in RNA binding and have at least one RRM domain in combination with other RNA binding domains (top 10 are shown in table S2). The most structurally homologous protein owing to the RRM fold include proteins that are involved in varied functions such as biogenesis, folding, splicing, maturation, stability, regulation and degradation of RNA. Out of the top 10 hits, 3 of them homodimerize in order to function consisting 1YTY, 5TKZ, 2M9K and many of them harbors both RNA binding and protein binding properties. These indicate the DND1 RRM2 dimer could be involved in protein interactions.

### DND1 RRM2 also forms closed monomeric structure

Structural analysis of the monomer was also performed in order to address whether the monomer had a closed structure where β1 and β4 are folded on to itself or an open structure as observed in the crystal structure described in this study. Sequence-specific backbone resonance assignment of the RRM2 domain was achieved using standard triple-resonance 3D NMR spectra. 2D [^15^N,^1^H] HSQC spectrum with labeled assigned peaks is shown in figure S2 A. 3D ^15^N-edited [^1^H,^1^H]-NOESY (mixing time=120 ms) spectrum measured on *U*-^15^N labeled protein was used to examine the monomer structure. We observed across the strands *N,N*(*i,j*) crosspeaks between S24 on E97 and D26 on A95 in the NOESY spectrum conforming the closed monomeric conformation (figure S2 B). Backbone chemical shift based structure prediction of monomer structure using CS Rosetta revealed that it was indeed a closed structure and upon comparison with open structure single chain showed a RMSD of 2.04 Å shown in figure S2 C. For the monomer to open conformational flexibility should be observed in the loop region as they have hydrophobic amino acids which are interacting and it would require many interactions to disrupt which will require large change in free energy to cross large kinetic energy barrier to form a dimer. For this ^15^N labeled protein was purified and both monomer and dimer population were collected separately after size exclusion chromatography. 2D [^15^N,^1^H] HSQC spectra was acquired for both that provided another evidence of existence of both monomer and dimer in solution state. We also observed increased line-width of resonance in the 2D [^15^N,^1^H] HSQC spectrum for the dimer form as compared to monomer due to increase in the rotational correlation time of the dimer (figure 3A). The chemical shift perturbation (CSPs) calculated after comparison of the backbone amide chemical shifts revealed changes occur in β1, α2, β4 and the hinge loop connecting α2 and β4 with maximal change in S87 from where the loop starts. While the major CSPs are due to direct structural changes between the open and closed form structure, the loop perturbations observed between α1 and β2 could occur due to indirect effects in spatial changes in protein structure figure 3C.

**Figure 3.**
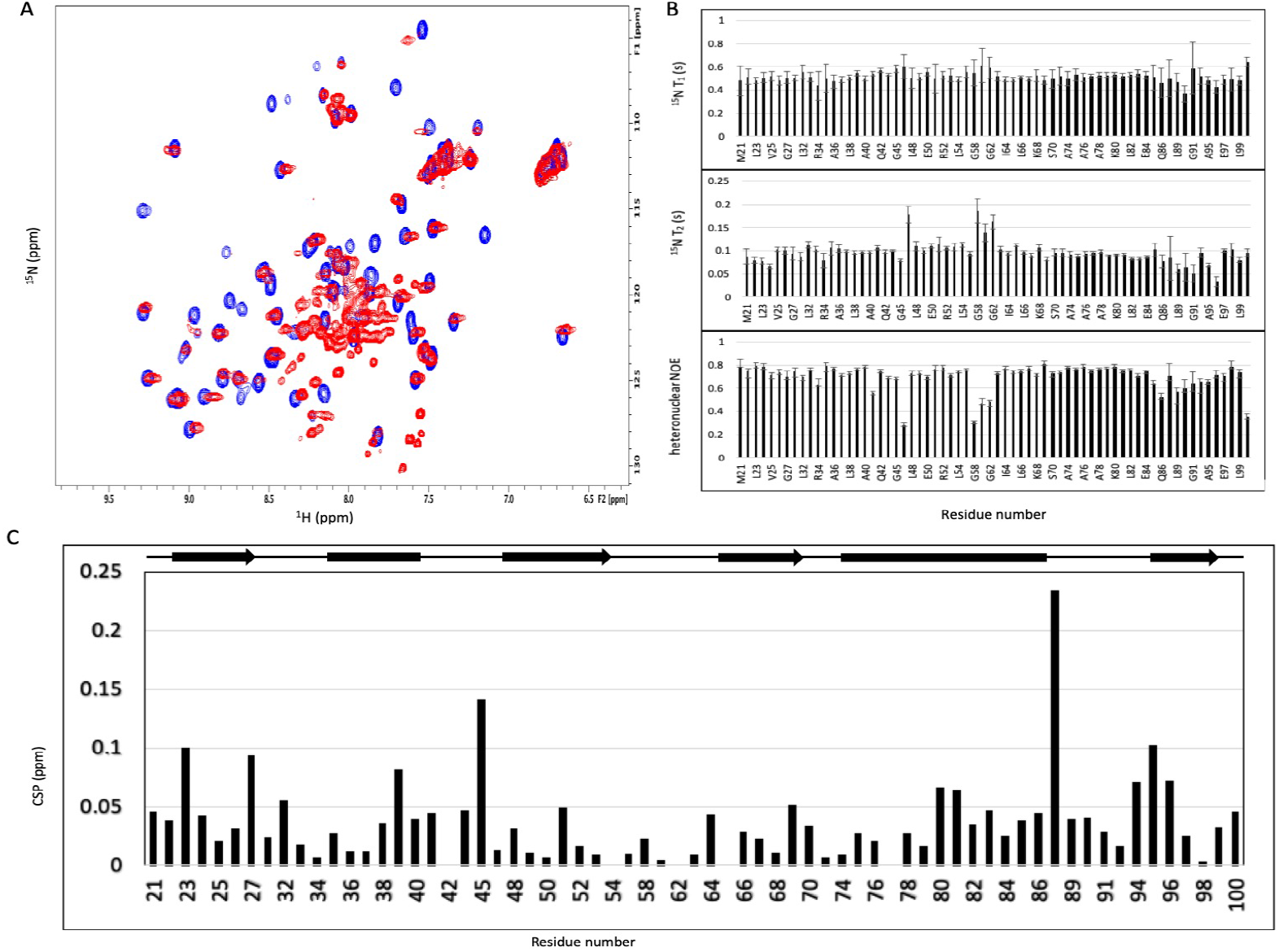
Solution state NMR characterization of conformational flexibility and dynamics. **(A)** Superposition of the 2D [^15^N,^1^H] HSQC spectra of the DND2 RRM2 in monomer and dimer states shown in blue and red, respectively **(B)** ^15^N backbone relaxation parameters T_1_, T_2_, ^15^N-{^1^H} nOe. **(C)** CSP between monomer and dimer is shown for each residue, the secondary structural arrangement of RRM2 is indicated as arrows (strand) and lines (helix) on the top.

^15^N-NMR relaxation parameters T_1_, T_2_ and heteronuclear ^15^N-{^1^H} nOe were measured for RRM2 domain. Relaxation measurement results are shown in figure 3b where the measured values are plotted against residue number. The values of the relaxation measurements (^15^N-{^1^H} nOe, T_1_, and T_2_) correspond to the mobility of each amide in the protein backbone. The average values for T_1_ and T_2_ were found to be 516 ± 56 ms and 96 ± 83 ms respectively shown in figure 3B. Estimated from T_1_/T_2_, the overall rotational correlation time (τ_c_) for the domain was 7.9 ns. The global correlation time estimation establish that the domain is monomeric under the experimental conditions used. Model Free analysis was performed based on the NMR relaxation parameters for each individual ^15^N providing motional information in terms of order parameters S2, internal correlation times τ e and contributions to chemical exchange Rex, for individual NH vectors across the domain backbone. The average S2 value across the structured region (residues M21–V83) was 0.90 ± 0.23 which confirm completely rigid nature of the domain. Although the secondary structure elements showed less variation, the residues belonging to the loop region showed elevated T_2_ implying conformational exchange. Loop 2 (L41-L48), loop3 (P55-G62) and loop 5 (S87-V94) shows faster motions with ^15^N-{^1^H} heteronuclear nOe values reaching as low as 0.3.

### RRM2 dimer is more stable than open monomer

Molecular dynamics simulations were performed for both open monomer and dimer for 1000 ns and 200 ns respectively with His-tag coordinates deleted from the structure in order to reduce unwanted fluctuations. The goal of these simulations was to account for protein flexibility, mobility and degree of rotation in the loop between α2 and β4, in case of open monomer and dimer and whether the monomer c terminal folds back on to itself to attain the conserved RRM fold. The RMSD of two trajectories is shown in Figure 4. The RMSD-Time evolution (Figure 4 A-B) shows that both the structures became stable after 100 ns. But the RMSD value was quite high around 7.2 Å in the case of open monomer while it was around 3.6 Å in the case of dimer. This indicates that the dimer conformation was much more stable compared to open monomer conformation. The root mean square fluctuation (RMSF) values were computed for each residue in DND1 RRM2 domain to recognize flexible and constrain region of the protein useful for characterizing local changes along the protein chain. The local changes in these residues indicates their importance during interactions. Residues which are more flexible usually have higher RMSF compared to constrain region with lower RMSF (Figure 4C and 4D). RMSF value of each residue in DND1 fluctuates maximally in the loop region between α1 and β2 (L41-G47), β2 and β3 (P55-G62) and α2 and β4 (H88-V94). While the α helix and β strands showed minimal fluctuations and were more rigid.

**Figure 4.**
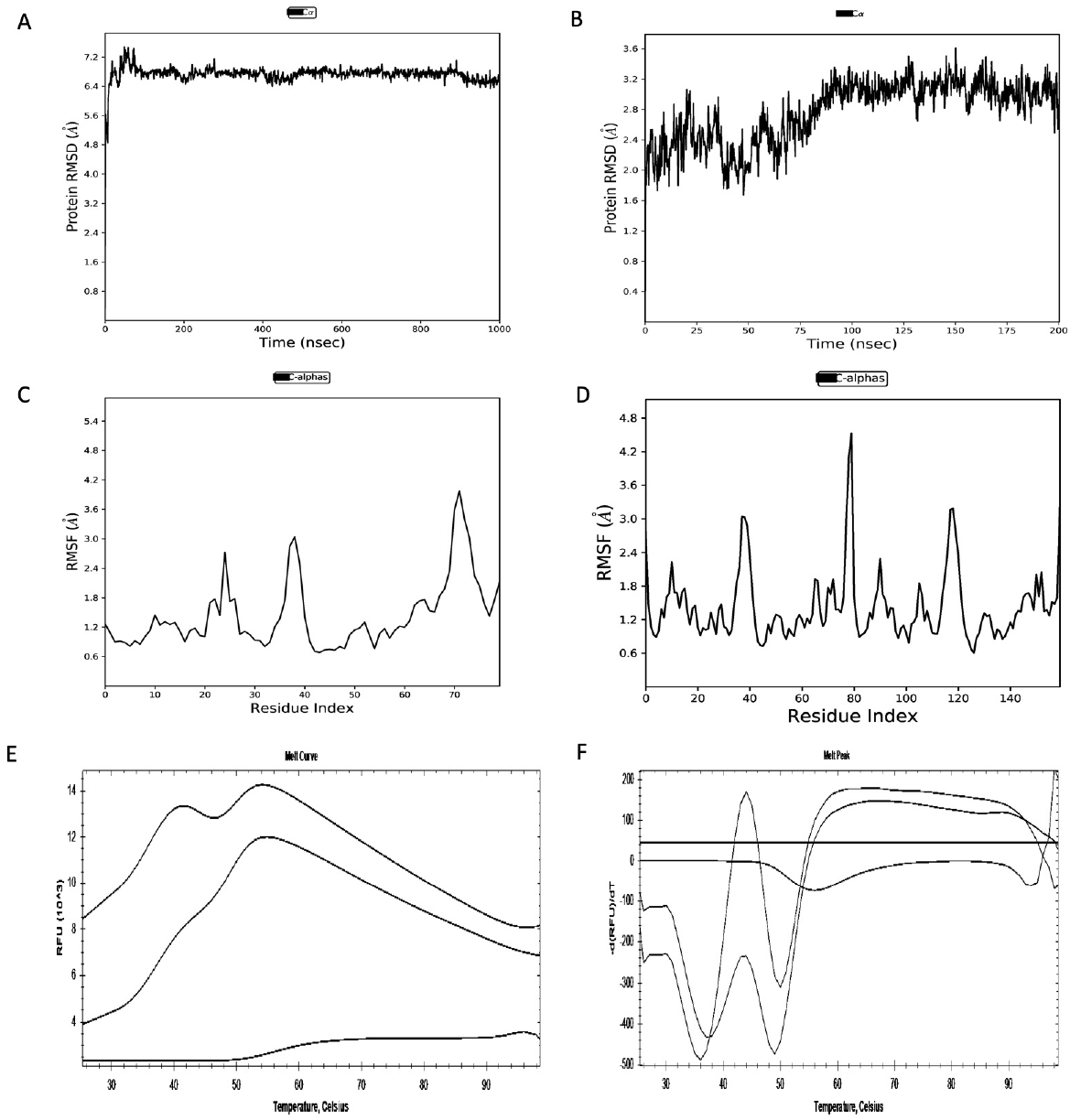
Concentration based Monomer-dimer switch. Root-mean-square deviation (RMSD) vs simulation time curves for two MD systems **(A)** open monomer for 1000 ns **(B)** Dimer for 200 ns. Protein root mean square fluctuation (RMSF) values along each residue in DND1 RRM2 domain **(C)** Monomer having 80 residues starting from M21 **(D)** Dimer having 160 residues. Thermal shift assay of DND1 RRM2 protein showing **(E)** melt curve and **(F)** melt peak at different concentrations.

The thermal shift assay, also called differential scanning fluorimetry, is a fast and inexpensive assay used for purified recombinant proteins to identify stabilizing buffer conditions, additives and small molecule ligands. It is a screening technique which also guides us to identify the suitable buffer pH in which the recombinant protein is most stable which further helps during crystallization trails. Here, we have used it to find out the protein melting temperature by using an external environment sensitive probe, and the fluorescence readout increases as the protein unfolds (Huynh and Partch, 2015). The protein was incrementally heated from 25 °C to 99 °C and as it unfolds, the hydrophobic core is exposed providing nonpolar region for the fluorescent dye to interact. Here we show in figure 4 E-F the melting curve of protein in different concentration harboring both populations in different ratios i.e., monomer and dimer. The concentration of recombinant protein used were 0.5mM, 1mM, 2mM shows one melting peak at 0.5mM while two peaks are observed in case of 1mM, 2mM conc. Having high dimer population in 2mM compared to high monomer population in 1mM. The thermal denaturation profile indicates a well-folded protein with low initial fluorescence at room temperature and as the temperature increases it leads to cooperative unfolding with high fluorescence giving rise to sigmoid curve. Also, the dimer unravels first as it is supported by fewer bonds when compared to monomer which is shown in the structure. The observed Tm here is 55 °C, which indicates that the overall domain is quite stable.

## Discussion

In order to understand the disparity in molecular function of DND1 and its role in development and cancer, which intrigued us to comprehend the mechanism of its regulation in cell, we determined the crystal structure of the DND1 RRM2 domain at 2.3 Å resolution. The structure of DND1-RRM2 reveals that the canonical RRM fold is being maintained by formation of a domain swapped dimer and the conformational flexibility is attributed due to the hinge α_2_-β_4_ loop that helps in mediating a domain swap forming anti-parallel β-sheet.

Residue level dynamics probed by solution-state NMR spectroscopy and MD simulations have also provided additional evidences for stable dimer formation. This is the first report of domain-swapped dimer formation in an RRM.

Evolution of protein oligomers is favored as it possesses several advantages over their monomers and 3D domain swapping is one such mechanism by which oligomerization is achieved. Protein oligomerization provided a larger binding surface, possibility of allosteric control and formation of active sites that has many biological implications. It also acts as a mechanism for regulating protein function, and as an evolutionary strategy to create protein interaction network and molecular machines (Liu and Eisenberg, 2002). The dynamic switch between monomer to dimer can aid in executing a function.

A possible mechanism for 3D domain swapping is described where temporary conditions such as high protein concentration or change in pH allows a closed monomer to open (Bennett, et al., 1995). During the process it causes perturbation of the existing bonds leading to displacement of a connecting loop as a free segment. The open monomer is then stabilized by formation of the same bonds with a swap domain with one or more subunits. For this switch to happen various types of interactions are initially broken and formed at the same protein interfaces, which includes hydrogen-bonds, electrostatic interactions, hydrophobic interactions, and disulfide bridge interactions. Individually, both the monomer and dimer are stable but the monomer requires activation energy to convert and form domain swap dimer. The temporary conditions which allows the bonds to breaks at the closed interface in monomer provides this energy. Therefore, activation energy for 3D domain swapping varies among domain-swapped proteins with diverse sizes and sequences of the swapped domains. Although domain swapping is relatively rare, several protein have been reported which functions as domain swapped dimer which include dipthera toxin, Bovine serum RNase, RNase A, CksHs2, SH2, SH3, staphylococcal nuclease, chymotrypsin inhibitor 2 etc. which have been extensively reviewed in several studies (Bennett, et al., 1995; Rousseau et al., 2003).

We have observed the RMSD between the closed and open monomers of DND1 RRM2 is 2.04 Å, which is quite large indicating high conformational flexibility. The conformational flexibility depends upon hinge loop length, hinge loop residues, solvent exposed surface area. Experimental evidence suggests that the length of the hinge loop if reduced leads to a more favored dimer conformation as the shorted loop is unable to fold back on itself and attain an open conformation more favored to domain swap on a contrary note very long loop is also disfavor domain swapping as it allows higher conformational plasticity (Nandwani et al. 2019; Ogihara et al., 2001). In the case of DND1 RRM2, 8 residues long hinge loop offers desired conformational flexibility to enable the transition between the closed monomeric structure and the open domain swapped dimer.

Recent evidence shows that introduction of a hydrophobic five residue motif comprising the sequence QVVAG in hinge loop leads to domain swapping irrespective of the position be it N terminal, C terminal or the middle of the protein (Nandwani et al. 2019). The hinge loop between α2 and β4 in DND1 RRM2 comprise of the surface exposed hydrophobic sequence GEQVAV, which is very similar and naturally present in the protein, which allows the domain swapping. The solvent exposed hydrophobic residues experience strain, which do not allow the loop to fold back upon itself similar to our observation in DND1 RRM2 having all hydrophobic amino acid in solvent exposed surface. The presence of glycine in the hinge loop region of RRM2 between α_2_ and β_4_ may provide considerable internal rotation and degree of flexibility. Glycine with the smallest cross-sectional area of about 4 sq Å has much more conformational flexibility in comparison to any other amino acid. Since, free rotation about -C(HR)-N(H)- bonds will be high for a hydrogen atom in comparison to any other R group. Experiments such as NMR dynamics and MD simulation support our claim of relatively higher dynamics in hinge loop attributing to domain swapped dimer formation, which is more stable in comparison to open monomer. Also, the buried surface area in this case is more in dimer when compared to open monomer, which demonstrate that the associated species are more stable than the individual units.

Redox sensing by a protein acts as molecular switch which allow intermolecular or intramolecular disulphide bond formation regulating functions of proteins such as enzymes, receptors, transcription factors etc. (Nagahara, 2011; Wouters et al.). Rat mercaptopyruvate sulfurtransferase also shows monomer dimer switch under such redox regulation (Nagahara, 2018). The hinge loop in RRM2 also consist of an exposed Sulphur atom in cysteine residue which when present in oxidizing condition favors this dimerization by forming a disulphide bond. On the contrary, under strong reducing condition the extent of dimerization was reduced. This redox switch, not observed previously in any domain swapped RRM dimer, provides additional regulation to the transition in DND1 RRM2.

Previous studies have reported 3D domain swapping possibly linked to protein aggregation and misfolding (Jaskólski 2001). Recent studies show that there are several transcription factors such as Bax, FOXP3, SH2, SH3 which swap their domain for their function while in several this leads to misfolding and aggregation. Some of the domain swapped dimers are associated with disease or forming amyloids or amyloids like fibrils consist of stefins, cystatin, prion protein (Bennett et al., 2006). Although there are no reports where RRMs have formed domain swap structure, still a study has indicated formation of domain swap like intermediate state in TDP-43 RRM2, which forms cytoplasmic aggregates together with its C-terminal domain leading to sporadic amyotrophic lateral sclerosis (Mackness et al., 2014).

Several RRMs have been extensively studied to date and found to have versatile mode of interaction be it with nucleotides or proteins. RRM mediated protein-protein interactions have been categorized on the basis of the interaction surface (Muto and Yokoyama, 2012). This includes interaction surface consisting of α-helix, β-sheet and loop regions. Competitive interaction by cognate nucleotide and interacting protein is also observed. CFIm25/CFIm68 RRM/RNA complex is one such example where dimer association of CFIm25 and loop mediated interaction with CFIm68 is observed (Yang et al., 2011). While, further studies will be required to dissect the biological relevance of dimerization in the function of DND1. Collectively, our results show the major determinant which allowed 3D domain swapping in RRM2 protein, increasing the binding surface area which may have role in protein interaction rather than RNA recognition.

Our study supports a model of DND1 architecture that allows varied roles for its two RRM domains such as RNA as well as protein interaction. DND1 acts as an adapter protein with its RRM1 domain responsible for specific RNA recognition while RRM2 and dsRBD are putative domains, which could be responsible for protein-protein interactions. Our results provide a starting point for understanding how DND1 act as multifunctional adapter protein between mRNA and CCR4-NOT complex.

## METHOD DETAILS

### Protein domain prediction and Multiple Sequence alignment

DELTA-BLAST (Domain Enhanced Lookup Time Accelerated BLAST) (Boratyn et al., 2012) of Human DND1 was done and multiple sequence alignment was done using clustal omega (Sievers et al., 2011) from protein sequences from different phyla of vertebrates obtained from uniport consisting of Homo sapiens Q8IYX4, Mus musculus Q6VY05, Gallus gallus A0A1L1RQS8, Anolis carolinensis G1KMZ1, Xenopus laevis Q6DCB7, Danio rerio Q7T1H5.

### Cloning, overexpression and purification

Human *DND*1 (UniprotID Q8IYX4) coding sequence corresponding to RRM12 (56-217) was codon optimized for overexpression in *Escherichia coli* and chemically synthesized from GeneArt (Life Technologies). Sub-clones corresponding to the RRM1 (residues 56–136) and RRM2 (139–217) and the combined construct (56–217) were prepared. pET28a expression vector utilising *Nde*I and *Xho*I restriction sites, were used for subcloning all three constructs. The vector plasmid containing the desired construct was transformed in *Escherichia coli* BL21(DE3) *CodonPlus* cells and were used for protein overexpression having a thrombin-cleavable hexa histidine tag. The overexpression was done in *E. coli* BL21 (DE3) codon plus cells at 37/25 °C for 16-20 hours using 0.5 mM IPTG at A600 ~ 0.6 - 0.8. Purification of the overexpressed protein containing hexa-histidine tag was done by nickel-nitrilotriacetic acid (Ni-NTA) affinity resin, SP Sepharose cation exchange column followed by gel permeation chromatography on GE Healthcare 16/60 Hiload Superdex 75 in buffer 25 mM Tris-HCl (pH 8.0), 150 mM NaCl, 5 mM β-mercaptoethanol. L-Selenomethionine labeled protein was subsequently overproduced in selenomet base medium plus nutrient mix media supplemented with 40mg/l L-Selenomethionine. *U-*^15^N labeled and *U*-^15^N and ^13^C labeled-protein was prepared by over-expressing the protein in minimal M9 medium containing 1 g/L ^15^NH4Cl and 6 g/L glucose (for *U*-^15^N-labeled protein) or 1 g/L ^15^NH_4_Cl and 2.5 g/L ^13^C-glucose (for *U*-^15^N- and ^13^C-labeled protein).

### Crystallization, Data Collection and Structure Determination

Protein was concentrated to 30 mg/ml and crystallized by hanging drop method at 16 °C in Hampton crystal structure screen G2 condition 30% (w/v) PEG-MME 5000, 0.1M MES monohydrate pH-6.5, 0.2M ammonium sulfate and then with B4 condition of additive screen, crystals were flash-frozen after a brief soaking in mother liquor supplemented with 20% glycerol. X-ray diffraction data were collected at 100 K on beamline ID23A at a wavelength of 0.9795A with a Pilatus_6M_F detector at the ESRF-European Synchrotron Radiation Facility, Grenoble. Data were processed by autoPROC (Vonrhein et al., 2011). Structure was determined by the single wavelength anomalous diffraction (SAD) method using autosol and autobuild in Phenix (Adams et al., 2010). The model was refined by iterative cycles of manual model building in Coot (Emsley and Cowtan, 2004). Refinement progress was monitored by tracking the *R*_work_/*R*_free_ ratio (with *R*_free_ representing 5% of total reflections) (Kovalevskiy et al., 2018; Winn et al., 2011). Domain swapped structure was generated with PISA (Krissinel and Henrick, 2007). Crystallographic data collection and structure refinement statistics are presented in Table S1. Structural images were prepared with the PyMOL Molecular Graphics System, version 2.0 (Schrödinger, LLC).

### NMR spectroscopy

All NMR measurements were performed in 50mM sodium phosphate buffer pH 7.0 and 100 mM NaCl and 10 % (*v/v*) D_2_O measured on a 500 MHz Bruker Avance III spectrometer equipped with 5 mm TCI cryo-probe at 298K. Data were processed using Topspin 3.6 pl7 (Bruker). Sequence-specific backbone assignments of the protein were achieved using a set of standard double- and triple-resonance spectra (Bax and Grzesiek, 1993) namely 2D [^15^N,^1^H] HSQC, 2D [^13^C, ^1^H] HSQC, 3D HNCA, 3D HNCO, 3D HNCACO, 3D CBCACONH, 3D HNCACB, 3D CcccoNH, 3D HcccoNH, 2D [^13^C,^1^H]-HSQC [aliphatic (− 1 to 5 ppm ^1^H^ali^), aromatic (4.7 to 10 ppm ^1^H^aro^)] and 3D ^15^N [^1^H,^1^H]-NOESY (NOESY mixing time 120 ms). Backbone dynamics of the protein was measured using 2D ^15^N-{^1^H}-heteronuclear nOe and ^15^N *T*_1_, *T*_2_ relaxation experiments. Sixteen delays ranging from 10 to 1100 ms were used for *T*_1_, while 11 delays from 10 to 210 ms were used for *T*_2_. Protein backbone resonances were manually assigned using Computer Aided Resonance Assignment (CARA) (Keller, J., 2004) software with ^1^H, ^13^C, and ^15^N shifts referenced indirectly to the DSS methyl proton resonance at 0 ppm in all spectra. Chemical shift based monomer structure prediction was done using CS-Rosetta (Shen et al., 2008). Changes in chemical shifts (chemical shift perturbations, Δδ) of backbone amide protons were tracked on 2D [^15^N,^1^H] HSQC spectra recorded for monomer and dimer. CSPs were calculated using the equation

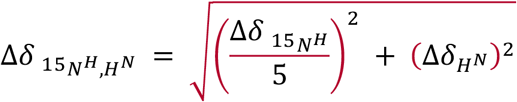

where Δ*δ*_H_^N^ and Δ*δ*^15^_N_^H^ are the changes in backbone amide chemical shifts for ^1^H^N^ and ^15^N resonances, respectively.

### Isothermal Titration Calorimetery

Protein was purified in buffer containing 25mM sodium phosphate buffer (pH 7.0) containing 100mM NaCl and 3mM β-mercaptoethanol. ITC experiments were performed with 0.1 mM protein inside the ITC cell and 1mM RNA in the syringe prepared in same buffer in a GE MicroCal iTC_200_ Calorimeter at 303K. 40 injections of 1 μl each were sequentially given containing RNA at 750 rpm mixing which gave heats that were further analyzed using Origin 7 software.

### Molecular Dynamics Simulation

For Molecular Dynamics (MD) simulations the Desmond program (Schrödinger, LLC) and OPLS3e (Harder et al., 2016) force field was used, with a simulation time set to 1000 ns for monomer and 200ns for dimer. For the simulations temperature was 300 K, pressure was 1.0325 bar, while cut off radius was set to 10 Å. The whole system was considered as isothermal–isobaric (NPT) ensemble class. TIP3P model (Berendsen et al., 1981) was used for modelling of the solvent. MD system consisted of one molecule of the protein placed into the cubic box. Input and output files in the case used were prepared on protein preparation wizard (Madhavi Sastry et al., 2013), analyzed and visualized with Maestro (Schrödinger, LLC) graphical user interface (GUI).

### Thermal Shift Assay

Melting temperature and concentration dependent dimer formation was tested using thermal shift assay for RRM2 protein. Increasing concentration of protein was taken and mixed with 5X SYPRO Orange dye and real time PCR run was setup initially at 25 °C, ramping up in increments of 1°C to a final temperature of 95 °C.

## DATA AND CODE AVAILABILITY

The atomic coordinates and structure factors (PDB ID: 6LE1) have been deposited in the Protein Data Bank (http://wwpdb.org). Partial backbone resonance assignment of DND1 RRM2 has been deposited in the Biological Magnetic Resonance Bank (BMRB) with accession number 50151.

## ACKNOWLEDGMENT

The study was supported by the ICGEB core funds. P.K is a recipient of CSIR senior research fellowship. We thank Dr. Alexander Popov (ID23-1 beamline, ESRF) for help with X-ray diffraction data collection. X-ray diffraction data collection at ID23 structural biology beamline was facilitated by the European Synchrotron Radiation Facility (ESRF) Grenoble, France Access Program, which is supported by Department of Biotechnology, Government of India [BT/INF/22/SP22660/2017]. The authors wish to thank DBT and ICGEB, New Delhi for providing financial support for the high field NMR spectrometers at the ICGEB, New Delhi and NII, New Delhi. The Department of Biotechnology, Government of India funded the ITC instrument through grant number BT/PR13018/BRB/10/731/2009 to N.S.B.

## AUTHOR CONTRIBUTIONS

P.K., and N.S.B. designed the experiments. P.K. performed the experiments. P.K., and N.S.B. analysed the data and wrote the manuscript. N.S.B. supervised the whole project.

## DECLARATION OF INTERESTS

The authors declare no competing interest.

