## supplementary data for "Human DND1-RRM2 forms a non-canonical domain swapped dimer"

**SUPPLEMENTAL INFORMATION**

**Table S1**. List of all the interactions seen between the two chains in the dimer calculated by PDBePISA (https://www.ebi.ac.uk/pdbe/pisa/)

| **No.** | **Chain:Residue** | **Residue no (atom)** | **Distance (Å)** | **Chain:Residue** | **Residue no (atom)** |
| --- | --- | --- | --- | --- | --- |
| 1 | A: GLU | 22 (O) | 2.94 | B: LEU | 99 (N) |
| 2 | A: SER | 24 (O) | 2.76 | B: GLU | 97 (N) |
| 3 | A: ASP | 26 (O) | 2.79 | B: ALA | 95 (N) |
| 4 | A: GLY | 27 (O) | 2.27 | B: GLN | 93 (NE2) |
| 5 | A: ASP | 26 (N) | 2.67 | B: ALA | 95 (O) |
| 6 | A: SER | 24 (N) | 2.76 | B: GLU | 97 (O) |
| 7 | A: CYS | 90 (SG) | 1.95 | B: CYS | 90 (SG) |

**Table S2**. DALI results. A. Table showing top ten results from a DALI search (http://ekhidna2.biocenter.helsinki.fi/dali/) of the DND1 RRM2 domain against the PDB dataset

| **No.** | **PDB ID** | **Z-score** | **RMSD** | **Name** | **Function** | **Annotation** |
| --- | --- | --- | --- | --- | --- | --- |
| 1 | 4eyt-C | 7.6 | 2.1 | p65 | Tetrahymena telomerase associated protein p65, holoenzyme of La family protein | RNA binding |
| 2 | 1yty-B | 7.6 | 3.1 | La NTD | Human La NTD protein-RNA complex | RNA binding |
| 3 | 2von-A | 7.5 | 2.5 | La NTD | Human La protein | RNA binding |
| 4 | 5tkz-B | 7.5 | 2.7 | MEC-8 | *C. elegans* RBMPS family protein | DNA binding |
| 5 | 4n0t-A | 7.5 | 2.4 | U6 snRNP | *S. cerevisiae* U6 small nuclear ribonucleoprotein | RNA binding |
| 6 | 6gx6-A | 7.3 | 3.5 | IMP3 | Human insulin-like growth factor 2 | RNA binding |
| 7 | 2m9k-A | 7.3 | 3.0 | RBMPS2 | Human RBMPS2 protein | RNA binding |
| 8 | 6e4n-A | 7.2 | 2.0 | TbRGG2 | *T. brucei* Kinetoplastid RNA (kRNA) editing  protein | RNA binding |
| 9 | 5lsb-F | 7.2 | 1.6 | Hsh49p | *S. cerevisiae* U2 snRNA SF3b component | RNA binding |
| 10 | 5d78-A | 7.2 | 2.5 | Mip6 | *S. cerevisiae* RRM3 of Mip6 | RNA binding |

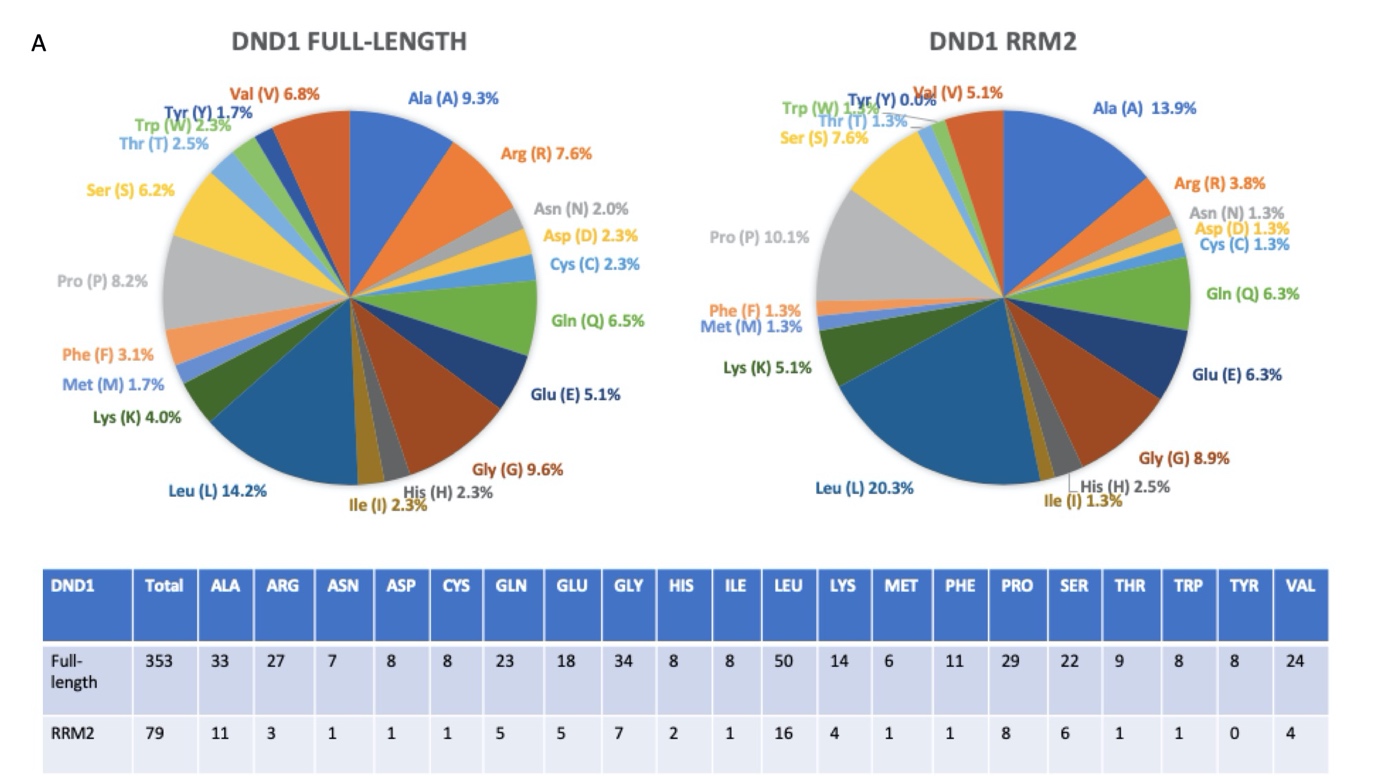

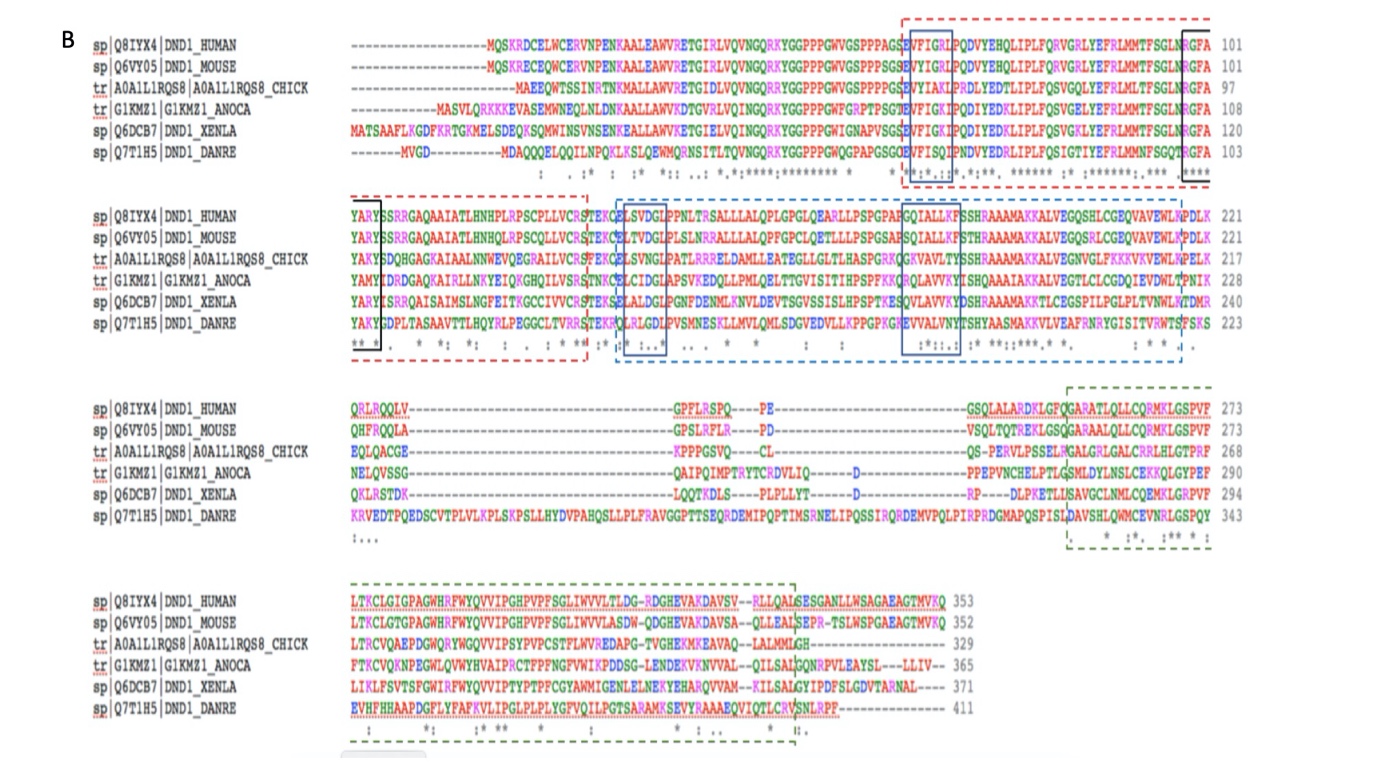

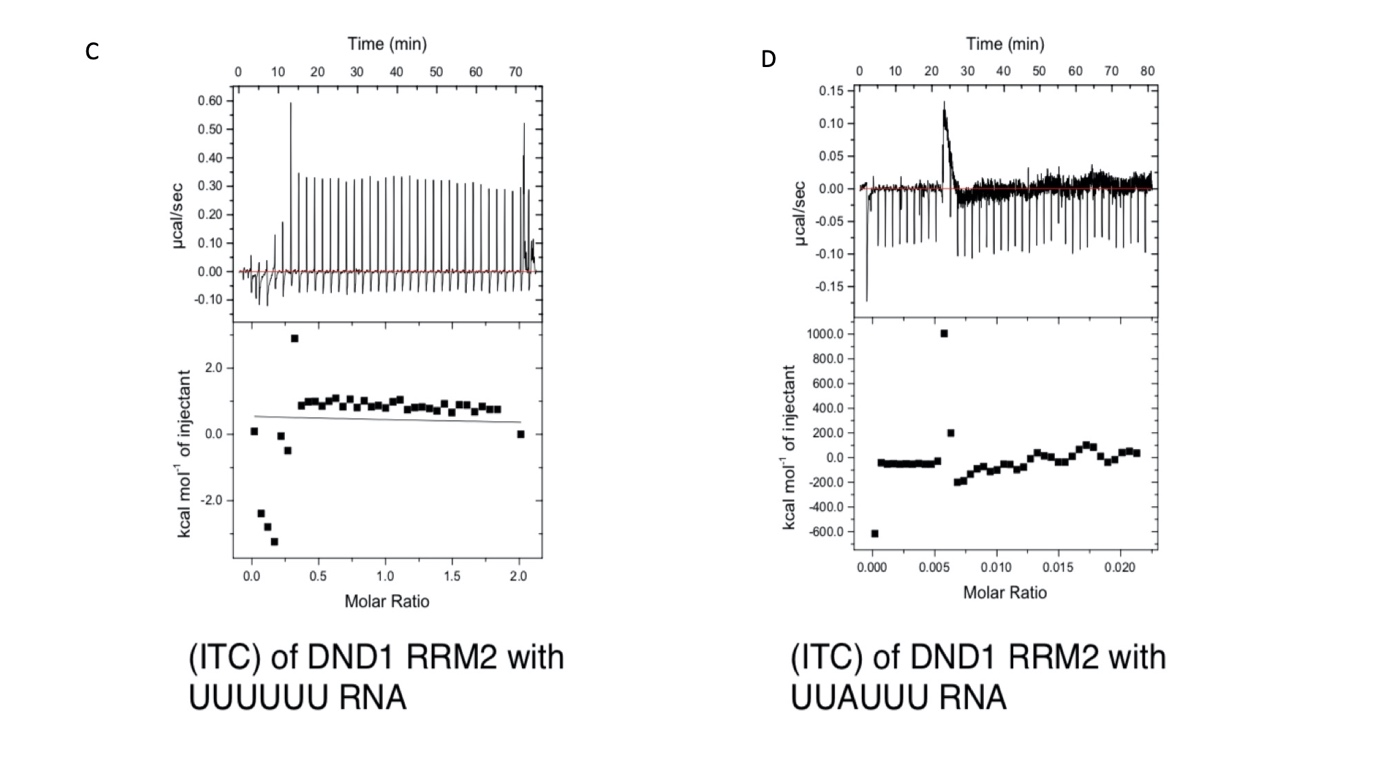

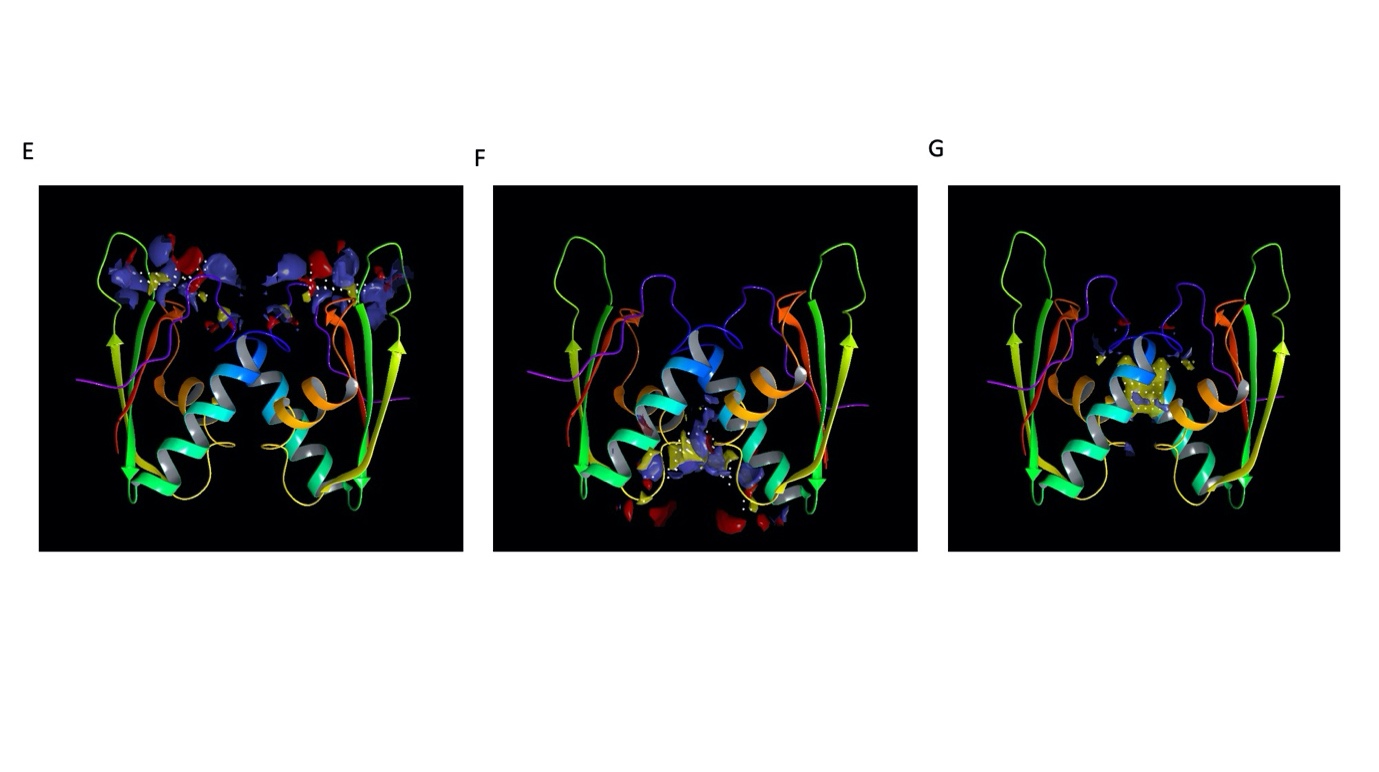

**Figure S1 Amino acid abundance, sequence conservation and associated functional significance of RRM2** **(A**) full length DND1 and DND1-RRM2 showing significant bias consisting 50% residues, which are aliphatic having distinctly high content of leucine, alanine, glycine and proline, 7% aromatic, 20% neutral, 13% basic and 7% acidic residues.

**(B)** Sequence alignment of DND1 protein sequence from different classes in vertbrates consisting of fish, amphibian, reptile, bird and mammals consisting of representative species Zebrafish (*Danio rerio* Uniprot ID- Q7T1H5), Chiken (*Gallus gallus* Uniprot ID- A0A1L1RQS8), Chameleon (*Anolis carolinensis* Uniprot ID- G1KMZ1), frog (*Xenopus laevis* Uniprot ID- Q6DCB7), Mouse (*Mus musculus* Uniprot ID- Q6VY05), Human (*Homo sapiens sapiens* Uniprot ID- Q8IYX4) showing RRM1, RRM2 and dsRBD boundaries, RNP sites and their conservation.

**(C)** Isothermal Titration Calorimetry of RRM-RNA binding RRM2 protein titrated against UUUUUU **(D)** RRM2 protein titrated against UUAUUU showing no interaction upon titration as the RNP sites containing the aromatic residues which bind RNA are not well conserved.

**(E), (F), (G)**. Binding surface identification using sitemap showing regions in the protein acting as hydrogen bond donor shown in blue, hydrogen bond acceptor shown in red and hydrophobic region shown in yellow.

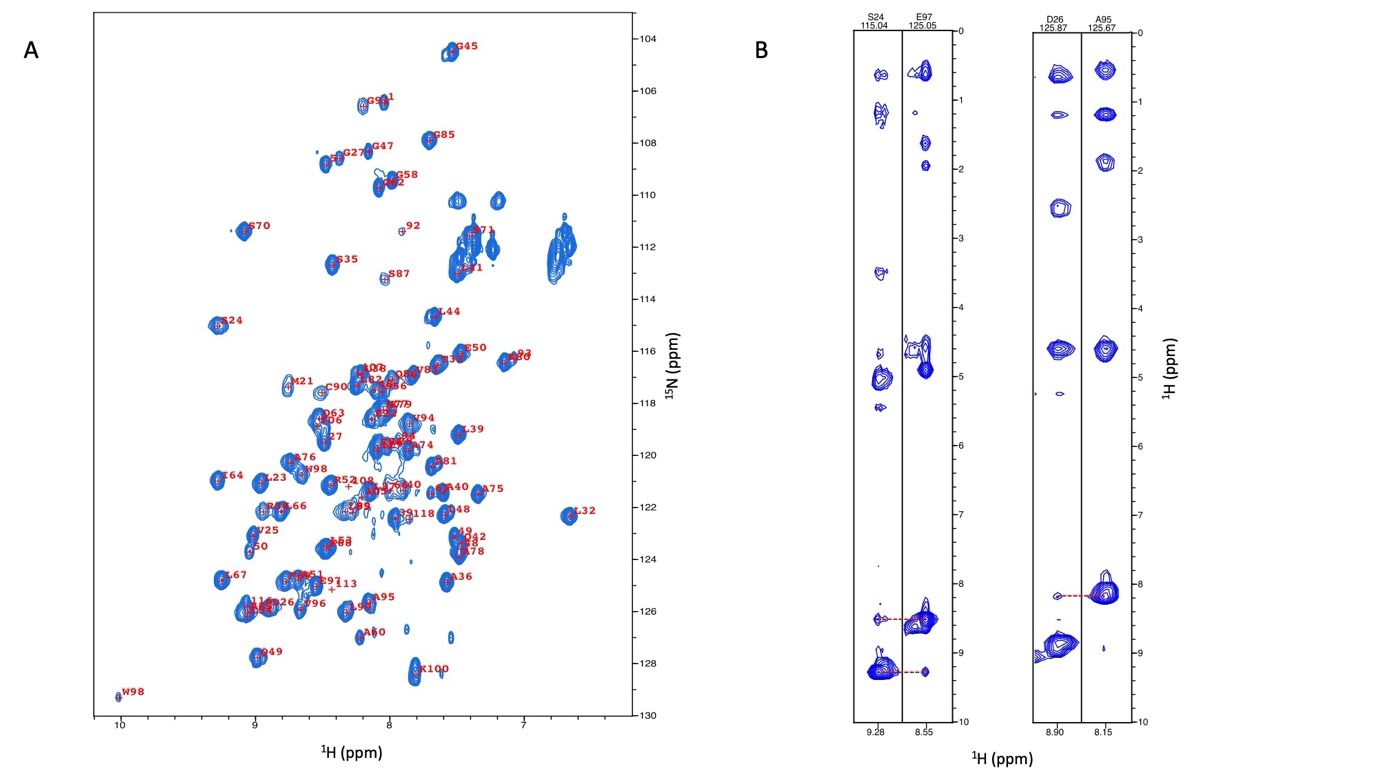

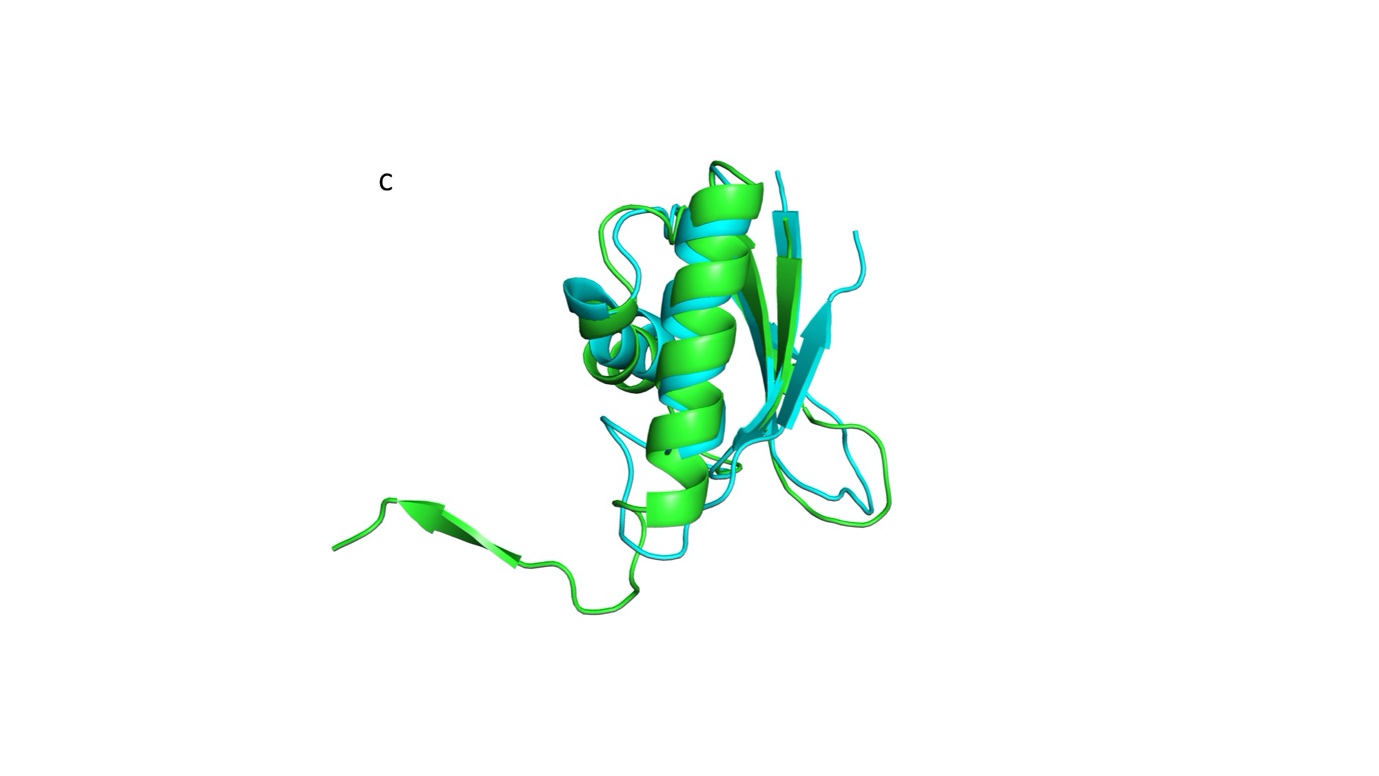

**Figure S2** **Backbone resonance assignment of RRM2 and structure comparision**

**(A)** 2D [^15^N,^1^H] HSQC spectrum of DND1 RRM2 with backbone ^15^N and ^1^H^N^ resonance assignments labeled **(B)** Strips from 3D ^15^N [^1^H,^1^H]-NOESY spectra showing long-range backbone ^1^H^N^-^1^H^N^ nOe connectivities between S24-E97 and D26-A95 across the β_1_ and β_4_ strands in the closed conformation of the DND1 RRM2 monomer. nOe connectivities are marked in red dotted line

**(C**)RMSD 2.04 Å upon morphing of closed monomer (cyan) and open monomer (green) DND1-RRM2 structures.
